# TRAPPC8 Is an Endogenous Brake on UFMylation That Suppresses Tauopathy

**DOI:** 10.64898/2026.08.18.745347

**Authors:** Yuansong Wan, Ethan Cordes, Weixi Feng, Se-In Lee, Katie Munechika, Yu Sun, Zhisong Gao, Beichen Gao, Jingjie Zhu, Man Ying Wong, Kendra Norman, Siyu Wang, Hao Chen, Bangyan Liu, Zhong Li, Mahalashmi Srinivasan, Sadaf Amin, Xiaomu Wei, Sue-Ann Mok, Rong Shen, Wenjie Luo, Shiaoching Gong, Hongmin Li, Haiyuan Yu, Li Gan

## Abstract

UFMylation, a ubiquitin-like protein modification, drives tau spread through the brain, but what keeps this process in check has remained unclear. We show that TRAPPC8 acts as a natural brake on UFMylation, binding directly to the E1 enzyme UBA5 to dampen pathway activity. In Alzheimer’s disease brain tissue, TRAPPC8 is reduced while UFMylation is elevated, suggesting this brake fails as disease progresses. Sustaining UFMylation in human iPSC-derived neurons increased tau aggregation and spread while disrupting lysosomal function and lipid balance. Restoring TRAPPC8– UBA5 binding reversed these defects through the lysosomal protein CLN8, which restored lysosomal function and reduced tau pathology. Notably, expressing just the UBA5-binding region of TRAPPC8 was enough to suppress tau pathology in vivo, marking it as a promising therapeutic target.

## Main Text

UFMylation is a reversible ubiquitin-like post-translational modification mediated by conjugation of ubiquitin-fold modifier 1 (UFM1) to target proteins. In this pathway, UFM1 is processed by the cysteine proteases UFSP1/UFSP2, activated by the E1 enzyme UBA5, transferred to the E2 enzyme UFC1, and ligated to substrates by the E3 ligase complex UFL1–DDRGK1 (*1, 2*) (*3, 4*). Emerging evidence implicates UFMylation in diverse cellular processes, including DNA damage responses, neurodevelopment, antiviral immunity, telomere maintenance, and tumorigenesis (*5–11*). Despite its broad functional relevance, the mechanisms that regulate UFMylation activity—particularly in neurons—remain poorly defined.

Recent studies suggest that dysregulation of UFMylation contributes to neurodegenerative disease. We previously showed that suppression of UFMylation, via knockdown of UBA5 or UFM1, attenuates tau propagation in human neurons(*12*). Consistent with this, alterations in UFMylation components have been reported in Alzheimer’s disease (AD) and amyotrophic lateral sclerosis(*13, 14*). However, the endogenous mechanisms that restrain UFMylation activity in neurons, and whether disruption of these regulatory pathways contributes to tau pathology, remain largely unknown.

Tauopathies are a heterogeneous group of neurodegenerative diseases characterized by the accumulation of pathological tau aggregates, including AD, the most common form, and frontotemporal lobar degeneration with tau pathology (FTLD-tau) subtypes such as Pick’s disease, corticobasal degeneration, and progressive supranuclear palsy (*15–17*). Pathological tau disrupts multiple cellular processes, including mitochondrial function, nuclear integrity, and organelle homeostasis(*18–21*) . Clinically, tau burden strongly correlates with cognitive decline and reduced survival in AD, underscoring tau as a central driver of disease progression and a key therapeutic target (*22–24*).

Here, we investigate the role and regulation of UFMylation in tauopathy. Analysis of postmortem AD brain tissue reveals elevated UFMylation in the cortex. We identify Trafficking Protein Particle Complex Subunit 8, TRAPPC8, as a previously unrecognized binding partner of UBA5 that functions as an endogenous brake on UFMylation. We further uncover a UFMylation–TRAPPC8–CLN8 axis that controls lysosomal integrity and lipid homeostasis. Mechanistically, sustained activation of UFMylation promotes tau propagation in human and mouse neurons, whereas increasing the UBA5-binding region (U5BR) of TRAPPC8 suppresses tauopathy both in vitro and in vivo. Downstream, restoring CLN8— Ceroid Lipofuscinosis, Neuronal 8—rescues lysosomal defects and attenuates tau aggregation, identifying it as a key effector of TRAPPC8-mediated protection. Together, these findings establish TRAPPC8 as an endogenous regulator of neuronal UFMylation and identify the UFMylation–TRAPPC8–CLN8 axis as a therapeutic pathway linking lysosomal homeostasis to tau pathology.

### Mapping UFMylation associated proteins in Alzheimer’s Disease Cortex and in human neurons

The transition from amyloid deposition to overt tau pathology marks a decisive step in AD progression (*25, 26*), yet the upstream proteostatic mechanisms that facilitate tau accumulation and spread remain incompletely defined. Given our prior evidence that suppression of the UFMylation cascade attenuates tau propagation (*12, 27*), we asked whether UFMylation is altered in human AD brain and whether it mechanistically contributes to tau pathology.

We examined postmortem dorsolateral prefrontal cortex (Brodmann area 9; BA9) and superior temporal gyrus (Brodmann area 22; BA22) from 35 neurologically normal controls and 50 AD cases (**Table S1**). Immunoblot analysis revealed a robust increase in UFM1-conjugated protein species in AD relative to controls in both cortical regions (**Fig. 1A, 1B, 1D**, **and 1F, Supplementary Fig. 1A and 1B**). Multiple high-molecular-weight (HMW) UFM1-positive bands were selectively elevated in AD tissue, whereas free UFM1 levels were not comparably changed (**Supplementary Fig. 1C and 1D**). As expected, HMW tau species were markedly increased in both BA9 and BA22 (**Fig. 1C** and **1E**), suggesting co-occurrence of elevated UFMylation with pathological tau accumulation in disease-relevant regions.

To define the molecular substrates underlying this shift, we performed UFM1 antibody–based co-immunoprecipitation followed by mass spectrometry (Co-IP–MS) in control and AD cortex (**Fig. 1G**). Comparative analysis revealed pronounced remodeling of the UFM1 interactome in AD (**Fig. 1H**).

Proteins with increased association to UFM1 included the E3 ligase adaptor DDRGK1, consistent with enhanced UFMylation activity in disease. Gene ontology analysis of AD-enriched interactors highlighted pathways linked to mTOR signaling, protein secretion, lipid metabolism, and the unfolded protein response (**Supplementary Fig. 1E**) (**Table S2**), implicating UFMylation in lysosomal–autophagic and broader proteostatic regulation. Although the canonical substrate RPL26 did not differ between control and AD brains, multiple ribosomal proteins, including RPL17, RPL37A, and RPS14, were significantly enriched, suggesting altered ribosomal engagement in AD (**Fig. 1H**) (**Table S3**).

We next mapped the UFMylation interactome in human iPSC-derived neurons. Expression of UFM1– GFP or the E1 enzyme UBA5–GFP (**Fig. 1I**), followed by LC–MS, identified robust and overlapping binding partners (**Fig. 1J-L**), consistent with recovery of the core UFMylation machinery. Interactors were enriched for cytoskeletal factors, with MYH9, along with MYH11 and MYH10, among the most prominent candidates (**Fig. 1J**). Additional proteins, including PSMA3, LIMA1, DBN1, MRPL49, UPF1, and ENTR1, further link UFMylation to actomyosin organization, proteasomal function, and mitochondrial and RNA regulatory pathways in human neurons (**Fig. 1K**) (**Table S4**). TRAPPC8 was detected in both UFM1 and UBA5 pulldowns (**Fig. 1L**), indicating its association with the UFMylation machinery. UBA5-specific interactors, including TXNDC5 and CALR, connect UFMylation to ER and protein folding processes. Co-immunoprecipitation confirmed interaction of MYH9 with UFM1 (**Fig. 1M**) and association of TRAPPC8 with both UBA5 and UFM1 (**Fig. 1N**).

**Figure 1.**
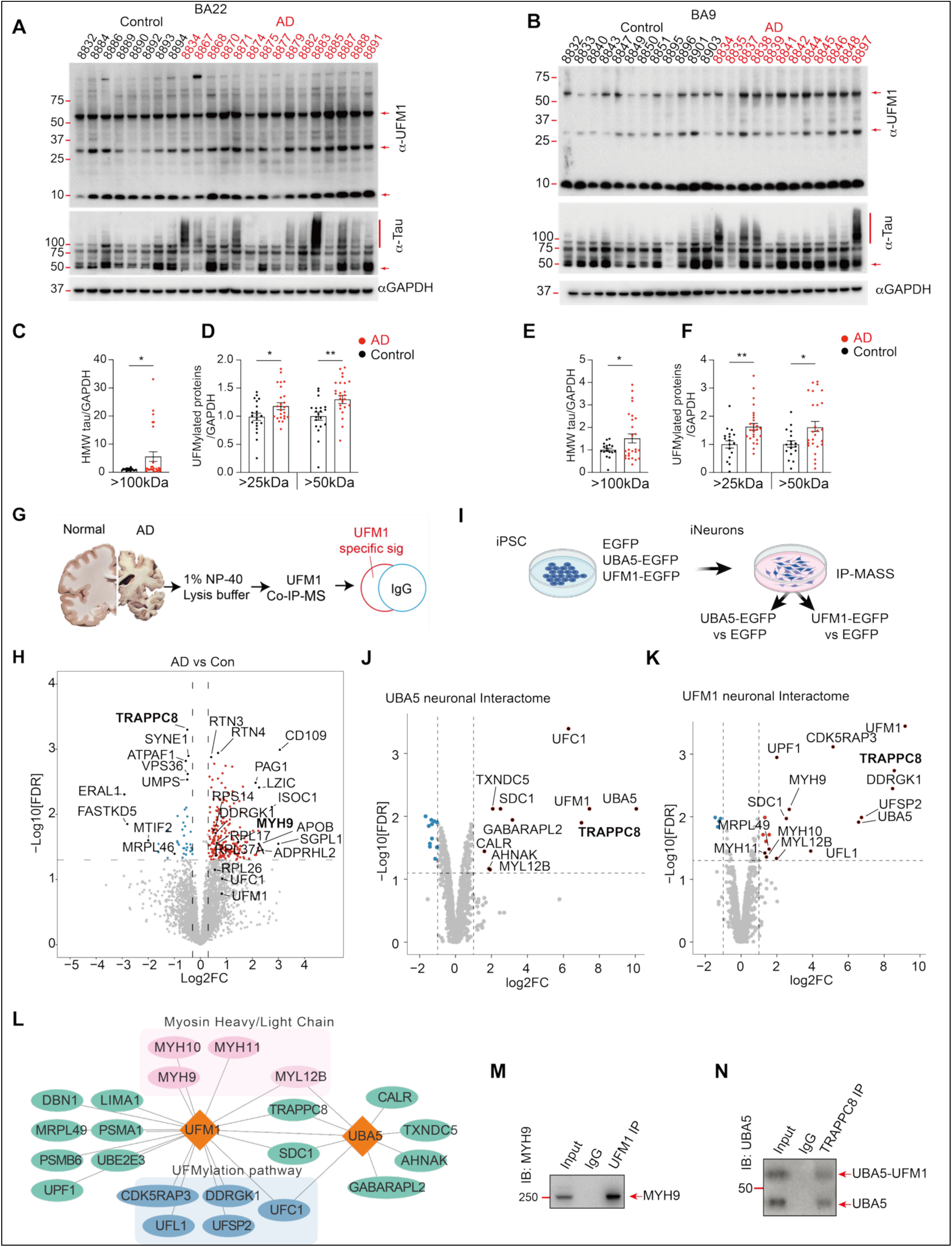
Mapping UFMylation associated proteins in Alzheimer’s Disease Cortex and in human neurons. (A, B) Representative immunoblots of UFM1 and tau in postmortem human posterior superior temporal gyrus (Brodmann area 22; BA22) (A) and dorsolateral prefrontal cortex (Brodmann area 9; BA9) (B) from neurologically normal controls and AD subjects. (C, E) Quantification of high-molecular-weight (HMW) tau species in control and AD samples from BA22 (C) and BA9 (E). (D, F) Quantification of UFM1-positive conjugated species, including upper bands (>50 kDa) and intermediate bands (>25 kDa), in BA22 (D) and BA9 (F). n = 20 (BA22), 17 (BA9) controls; n = 25 (BA22), 26(BA9) AD cases. Statistical significance was assessed by unpaired two-tailed Student’s t test (*p < 0.05, **p < 0.01). (G) Schematic of UFM1 immunoprecipitation followed by mass spectrometry (Co-IP–MS) to define the UFM1 interactome in control and AD cortex (N = 4 per group). (H) Volcano plot depicting differentially enriched UFM1-interacting proteins in AD relative to control brain. Dashed lines indicate thresholds of |log2 fold change| ≥ 0.5 and p < 0.05. (I) Diagram for UFM1-GFP and UBA5-GFP interactome study of human iPSC-derived neurons. (J, K) Volcano plot depicting differentially enriched UBA5-interacting proteins (J), and UFM1-interacting proteins (K) in iPSC neurons relative to EGFP controls. Dashed lines indicate thresholds of |log2 fold change| ≥ 1 and adjust p < 0.07 (J) and adjust p < 0.05 (K). n = 4 replicates from two independent experiments. (L) UFM1-GFP and UBA5-GFP neuronal interactomes identified by LC-MS and overlapped proteins interacting with both UFM1 and UBA5. (M, N) Endogenous co-immunoprecipitation to assay the binding of individual proteins in human iPSC-derived neurons. Protein lysates were immunoprecipitated for immunoprecipitated for UFM1 antibody and blotted for MYH9 antibody (M) or immunoprecipitated for TRAPPC8 antibody and blotted for UBA5 antibody (N).

### UFMylation depletes TRAPPC8; its restoration reverses proteome remodeling

To define how sustained UFMylation alters MYH9, TRAPPC8, and downstream proteostasis, we modeled chronic UFMylation activation in human neurons. Human iPSC-derived neurons expressing EGFP control, UBA5, or UFM1 were maintained for 21 days to induce persistent UFMylation. Immunoblotting confirmed robust accumulation of UFM1 conjugates upon UBA5 or UFM1 overexpression (**Supplementary Fig. 2A– C**). Under these conditions, TRAPPC8 protein levels were markedly reduced, whereas MYH9 shifted toward higher-molecular-weight (HMW) species (>250 kDa) with depletion of the canonical ∼250 kDa band (**Fig. 2A–C**). The enrichment of HMW MYH9 species is consistent with increased post-translational modification and supports MYH9 as a neuronal UFMylation substrate. To directly identify UFMylation sites on MYH9, we performed anti-UFM1 immunoaffinity enrichment followed by LC–MS/MS analysis. MS/MS spectra identified a UFM1-modified peptide containing the characteristic VG remnant on Lys1387 of MYH9, providing direct evidence that MYH9 undergoes UFMylation in neurons (**Supplementary Fig. 2D**). In contrast, TRAPPC8 did not exhibit a corresponding mobility shift but instead decreased in abundance, indicating it is unlikely to function as a substrate and instead acts as a regulator of the UFMylation pathway.

We next mapped how TRAPPC8 modulates UFMylation-driven proteome remodeling using quantitative proteomics comparing control, hyper-UFMylation, and hyper-UFMylation with TRAPPC8 restoration (**Fig. 2D**). Hyper-UFMylation induced broad proteomic changes, prominently affecting endolysosomal and lipid metabolic pathways (**Supplementary Fig. 2E**) (**Table S5**). TBC1D2 and CLN8—key regulators of autophagosome–lysosome fusion and lysosomal enzyme trafficking—were significantly reduced, whereas DGAT1, PTGES3, ARV1, and BCKDHB were increased, consistent with impaired lysosomal function and dysregulated neutral lipid metabolism(*28–33*). Restoration of full-length TRAPPC8 reversed these alterations, shifting protein expression toward baseline (**Supplementary Fig. 2F, and Fig. 2D, E**). Correlation analysis revealed an inverse relationship between UFMylation-induced changes and TRAPPC8-mediated rescue (R = -0.85) (**Fig. 2E**), indicating that TRAPPC8 counteracts UFMylation-driven proteome remodeling.

Among these targets, CLN8 emerged as a key effector. UBA5 or UFM1 overexpression significantly reduced CLN8 protein levels in human iPSC-derived neurons, (**Fig. 2F–G**). Notably, TRAPPC8 overexpression restored CLN8 levels under hyper-UFMylation conditions, consistent with our quantitative proteomics results (**Fig. 2H, I**). These results further support TRAPPC8 as a key regulator that restrains UFMylation-driven proteome remodeling, including depletion of CLN8, providing a mechanistic link between aberrant UFMylation and neuronal proteostatic stress.

**Figure 2.**
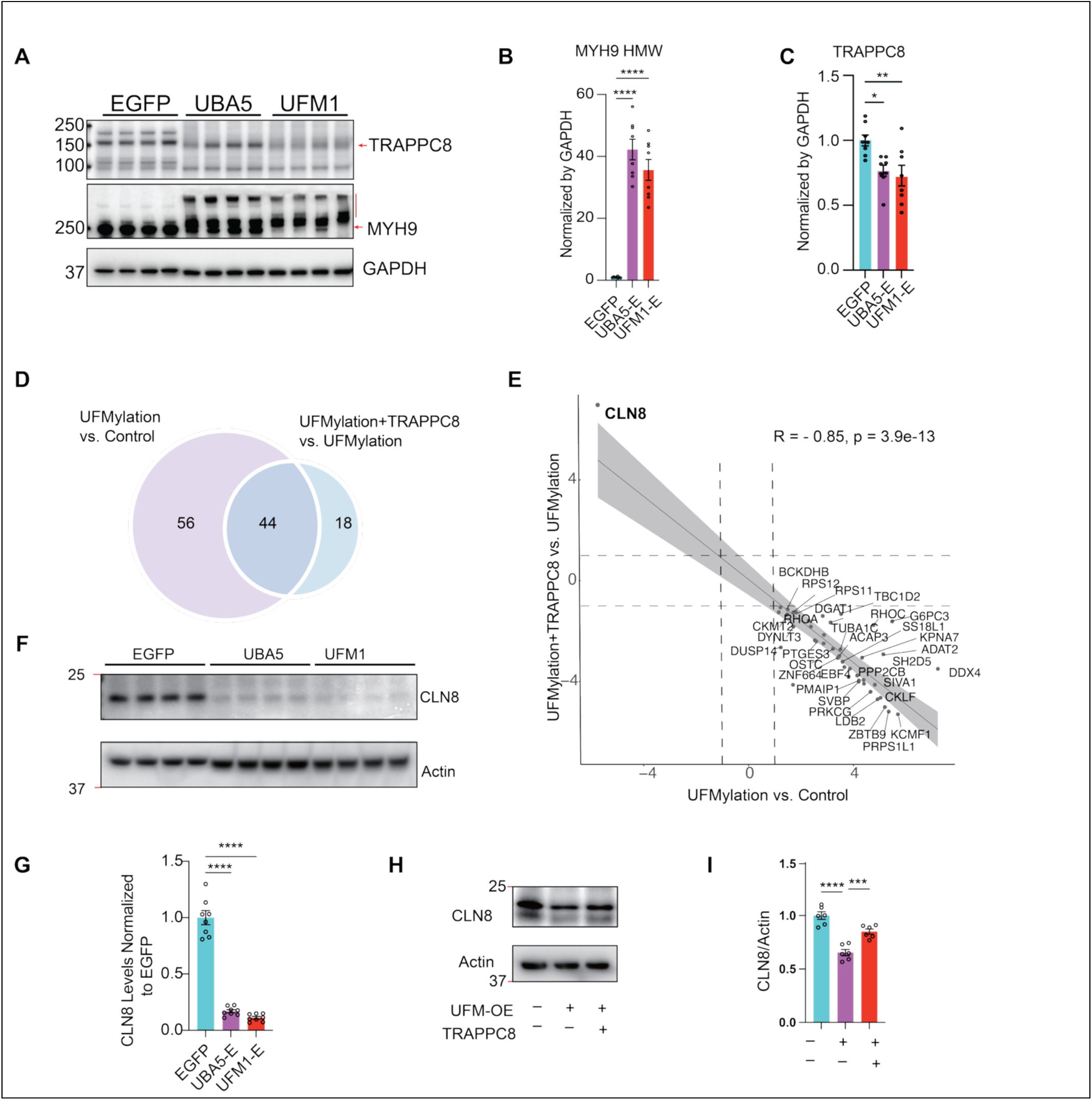
UFMylation depletes TRAPPC8 and overexpression of TRAPPC8 reverses UFMylation induced alterations. (A) Representative immunoblots of TRAPPC8 and MYH9 in human iPSC-derived neurons expressing EGFP control, UBA5, or UFM1 following 21 days in culture. (B, C) Quantification of TRAPPC8 (∼150 kDa), canonical MYH9 (∼250 kDa), and high-molecular-weight (HMW) MYH9 species (>250 kDa). n = 8 biological replicates from two independent experiments. Statistical significance was determined by one-way ANOVA with Tukey’s multiple comparisons test (****p < 0.0001). (D) Venn diagram illustrating differentially expressed proteins (DEPs) identified by proteomic analysis in UFMylation conditions versus control (left), and in UFMylation conditions with TRAPPC8 overexpression versus UFMylation alone (right). Overlapping proteins represent shared DEPs between the two comparisons. (E) Correlation analysis of overlapping DEPs from (D), comparing log2 fold changes in UFMylation versus control and UFMylation plus TRAPPC8 overexpression versus UFMylation alone. Dashed lines indicate |log2 fold change| ≥ 1. Pearson’s two-sided correlation test was applied. (F) Representative immunoblots of CLN8 and β-actin in human iPSC-derived neurons expressing EGFP, UBA5, or UFM1 after 21 days in culture. (G) Quantification of CLN8 protein levels in neurons. n = 8 biological replicates from two independent experiments. Statistical significance was assessed by one-way ANOVA with Tukey’s multiple comparisons test (****p < 0.0001). (H) Representative immunoblots of CLN8 and β-actin in HEK cells expressing EGFP, UBA5, or UFM1. (I) Quantification of CLN8 protein levels in HEK cells. n = 6 biological replicates from two independent experiments. Statistical significance was determined by one-way ANOVA with Tukey’s multiple comparisons test (****p < 0.0001)

### Hyper-UFMylation disrupts lipid homeostasis and promotes lipid droplet accumulation

Our results so far links UFMylation to lysosomal integrity and membrane trafficking via TRAPPC8–CLN8 axis. CLN8 is required for ER-to-Golgi transport of lysosomal enzymes and is essential for lipid trafficking and turnover(*34*), (*35*). We next directly examined whether hyper-UFMylation alters neuronal lipid homeostasis. Unbiased LC–MS/MS lipidomic profiling of DIV42 human iPSC-derived neurons revealed a significant increase in neutral lipid species, including triacylglycerols and sterol esters, under hyper-UFMylation conditions (**Supplementary Fig. 3A–C**) (**Table S6**). These changes are consistent with impaired lipid degradation and redistribution. To functionally challenge lipid handling capacity, neurons were exposed to oleic acid (OA), a monounsaturated fatty acid that promotes triglyceride synthesis and lipid droplet formation (*36*), in part by limiting autophagic flux. Neutral lipid staining demonstrated marked lipid droplet accumulation in neurons overexpressing UBA5 or UFM1, an effect that was further amplified by OA treatment (**Supplementary Fig. 3D–F**). Lipid droplets were detectable even in the absence of exogenous lipid supplementation after prolonged hyper-UFMylation in neurons 5 weeks in culture, whereas EGFP control neurons exhibited minimal lipid accumulation (**Supplementary Fig. 3G–I**). Consistent with enhanced lipid droplet biogenesis, expression of the lipid droplet-associated protein PLIN2 was significantly increased in hyper-UFMylated neurons (**Supplementary Fig. 3J–L**). These findings demonstrate profound disruption of lipid metabolism under sustained UFMylation activation.

### Loss of TRAPPC8 drives a feed-forward amplification of UFMylation

We next asked whether TRAPPC8 directly regulates UFMylation machineray. To test this, we depleted TRAPPC8 in human iPSC-derived neurons using shRNA and examined endogenous UFM1 conjugation. Neurons cultured for 10 days exhibited a marked increase in UFM1-conjugated species upon TRAPPC8 knockdown, whereas UFM1 knockdown reduced these species as expected (**Fig. 3A–F**). Quantification across multiple molecular weight ranges—including >25 kDa, ∼37 kDa, >50 kDa, and ∼150 kDa species— demonstrated a broad amplification of UFM1 conjugates in the absence of TRAPPC8. These data indicate that TRAPPC8 depletion is sufficient to enhance basal UFMylation in human neurons, positioning TRAPPC8 as a negative regulator of the pathway.

To determine whether this regulatory relationship extends to non-neuronal cells and general stress-responsive contexts, we turned to HEK cells. TRAPPC8 was disrupted using CRISPR/Cas9 and cells were acutely stimulated with anisomycin to activate stress signaling. Under these conditions, TRAPPC8 loss led to a robust increase in UFM1-conjugated species compared to control cells (**Fig. 3G–J**), consistent with enhanced UFMylation activity, with acute anisomycin stimulation further boosting the levels of a ∼37 kDa UFMylated protein. Conversely, overexpression of TRAPPC8 attenuated UFM1 conjugation both in presence and absence of anisomycin stimulations (**Fig. 3K–N**), supporting TRAPPC8 levels act as a rate-limiting determinant of UFMylation. Consistent with the notion that MYH9 is a UFMylation-sensitive substrate, TRAPPC8 knockout in HEK cells increased the abundance of high-molecular-weight MYH9 species while altering the balance between monomeric and polymeric forms (**Fig. 3O–Q**). Our results support a feed-forward vicious cycle, in which elevated UFMylation depletes TRAPPC8, and loss of TRAPPC8 further amplifies UFMylation and disrupts proteostatic balance.

**Figure 3.**
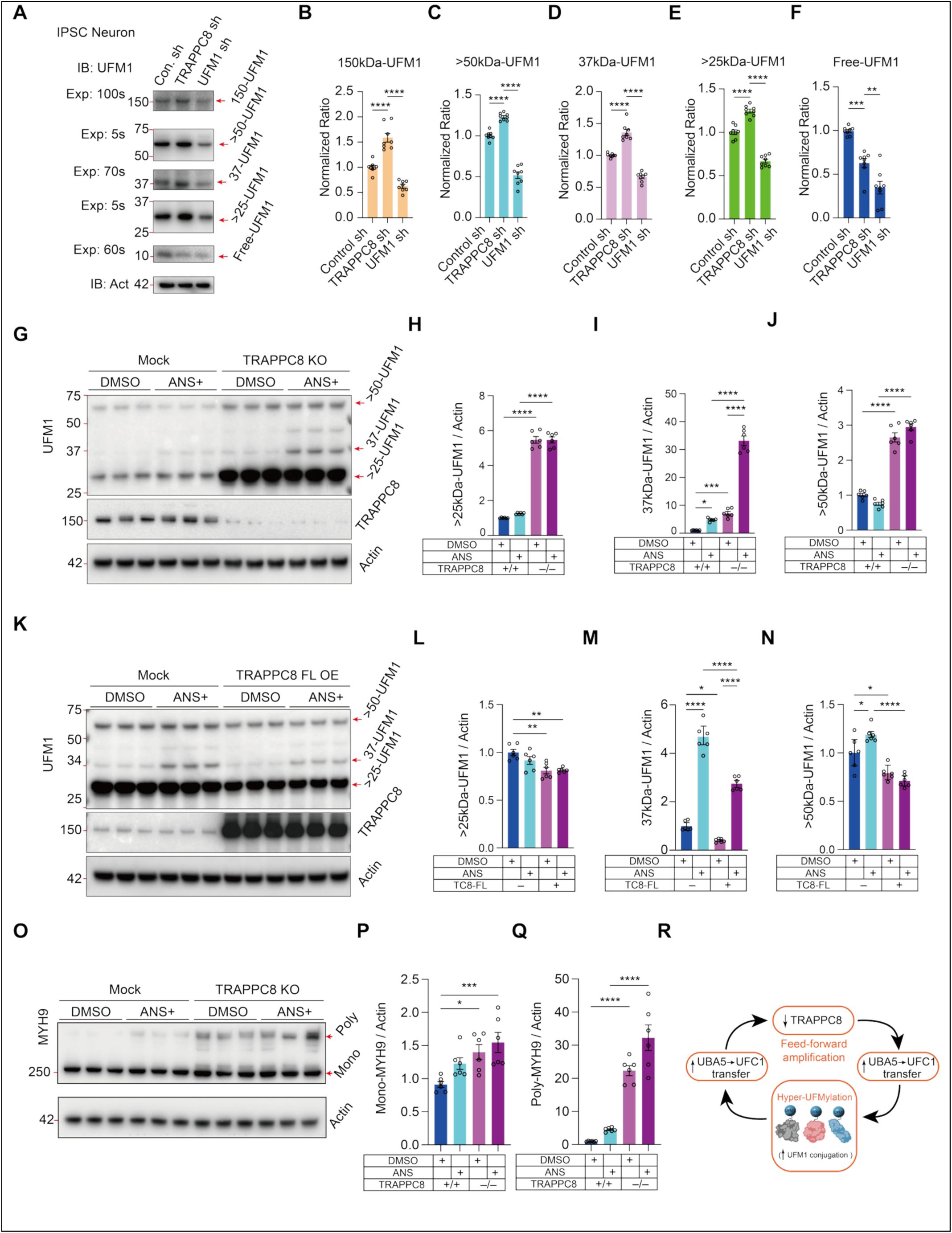
Loss of TRAPPC8 drives a feed-forward amplification of UFMylation. (A) Representative immunoblots of UFM1 and β-actin in human iPSC-derived neurons cultured for 10 days and transduced individually with control, TRAPPC8 shRNA, or UFM1 shRNA. (B–F) Quantification of UFM1-positive species, including the ∼10 kDa UFM1 monomer (B), >25 kDa (C), ∼37 kDa (D), >50 kDa (E), and ∼150 kDa (F) conjugated bands. n = 8 biological replicates from two independent experiments. Statistical significance was assessed by one-way ANOVA with Tukey’s multiple comparisons test (**p < 0.01, ***p < 0.001, ****p < 0.0001). (G) Representative immunoblots of UFM1, TRAPPC8, and β-actin in HEK cells transduced with control (mock) or TRAPPC8-targeting sgRNA cloned into lentiCRISPRv2. Cells were treated with 200 nM anisomycin (ANS) for 10 min prior to harvest. (H–J) Quantification of UFM1-conjugated species (>25 kDa, ∼37 kDa, and >50 kDa) in HEK cells described in (G). n = 6 biological replicates from two independent experiments. Statistical significance was determined by one-way ANOVA with Tukey’s multiple comparisons test (****p < 0.0001). (K) Representative immunoblots of UFM1, TRAPPC8, and β-actin in HEK cells transfected with control (mock) or TRAPPC8 overexpression plasmids, followed by 200 nM anisomycin treatment for 10 min. (L–N) Quantification of UFM1-conjugated species (>25 kDa, ∼37 kDa, and >50 kDa) in HEK cells described in (K). n = 6 biological replicates from two independent experiments. Statistical significance was assessed by one-way ANOVA with Tukey’s multiple comparisons test (****p < 0.0001). (O) Representative immunoblots of MYH9 and β-actin in HEK cells expressing control or TRAPPC8-targeting sgRNA, treated with 200 nM anisomycin. (P, Q) Quantification of monomeric and polymeric (high-molecular-weight) MYH9 species. n = 6 biological replicates from two independent experiments. Statistical significance was determined by one-way ANOVA with Tukey’s multiple comparisons test (*p < 0.05, ***p < 0.001, ****p < 0.0001). (R) Schematic model illustrating a reciprocal negative regulatory loop between UFMylation and TRAPPC8.

### TRAPPC8 loss phenocopies hyper-UFMylation to drive lysosomal dysfunction and tau aggregation

We next asked whether TRAPPC8 loss recapitulates the cellular consequences of hyper-UFMylation in human neurons. The downstream effects of elevated UFMylation on lysosomal and autophagic pathways were assessed in human iPSC-derived neurons expressing EGFP control, UBA5, or UFM1. After 21 days, immunoblot analysis showed that UBA5 or UFM1 overexpression reduced LAMP1 levels and decreased both LC3-I and LC3-II, consistent with impaired lysosomal and autophagic function (**Fig. 4A–C, Supplementary Fig. 4A–C**). To directly evaluate lysosomal integrity, we performed confocal imaging of lysosomal markers in neurons expressing EGFP, UBA5-EGFP, or UFM1-EGFP. In parallel, high-molecular-weight tau species (>70 kDa) were markedly increased (**Fig. 4D**). Hyper-UFMylation led to a pronounced reduction in lysosomal signal at both DIV16 and DIV35 (**Fig. 4E–F, Supplementary Fig. 4D–E**), corroborating the biochemical evidence of lysosomal deficiency. Moreover, UFM1 overexpression significantly increased MC1-positive inclusions (**Fig. 4G–H**), further supporting a prominent role of UFMylation in promoting tau accumulation and propagation.

TRAPPC8 deficiency is sufficient to reproduce these phenotypes. Knockdown of TRAPPC8 in human iPSC-derived neurons reduced LAMP1 levels and altered LC3 processing, closely mirroring the effects of UBA5 or UFM1 overexpression (**Fig. 4I–L**). Notably, TRAPPC8 depletion also increased MC1-positive tau aggregates (**Fig. 4M–N**), phenocopying the pro-aggregation effect of elevated UFMylation. Thus, loss of TRAPPC8 is sufficient to drive lysosomal dysfunction and enhance tau aggregation, recapitulating the cellular consequences of elevated UFMylation and reinforcing TRAPPC8 as a critical brake on this pathway.

**Figure 4.**
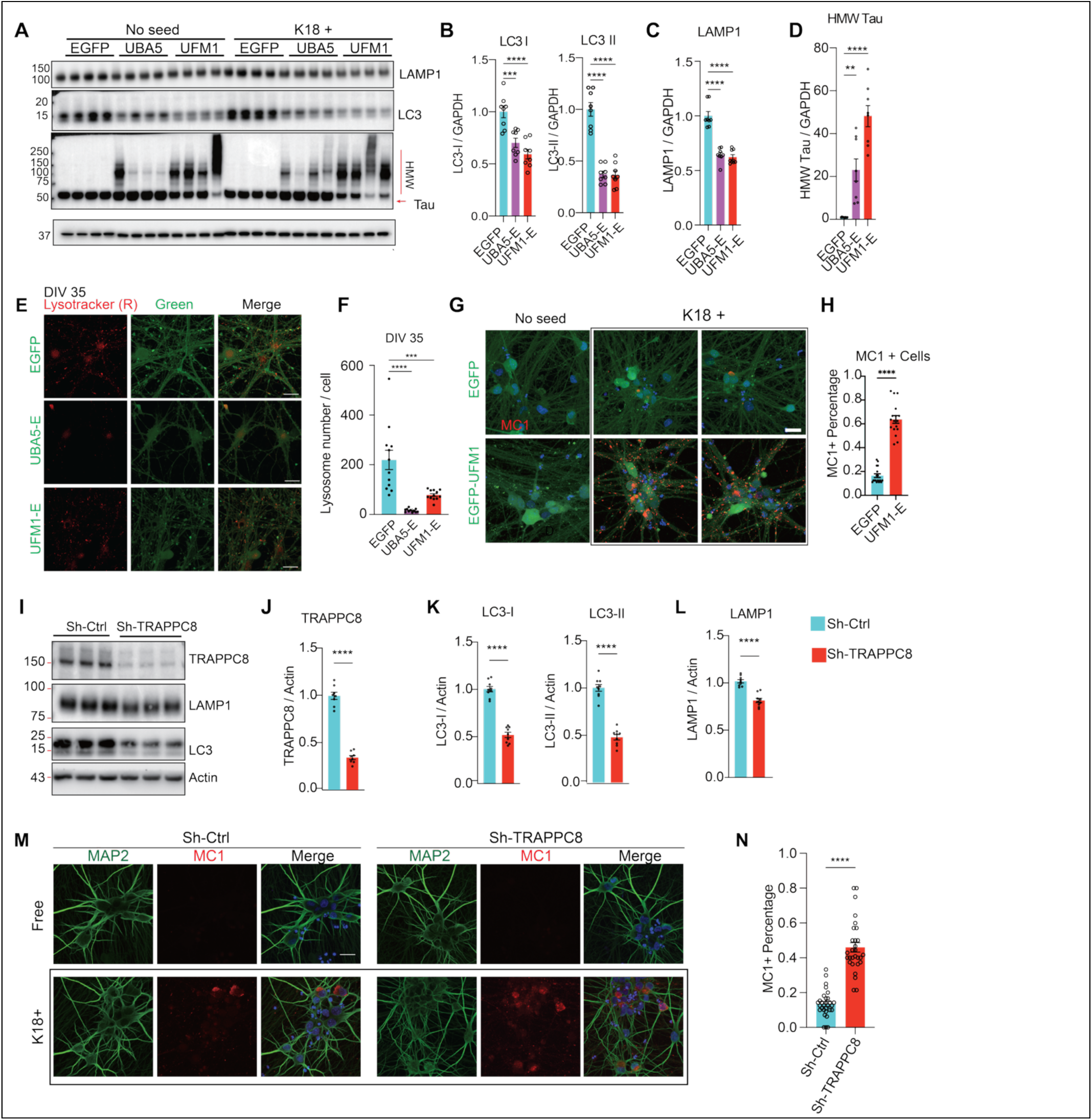
TRAPPC8 deficiency phenocopies UFMylation-induced lysosomal defects and enhances tau propagation. (A) Representative immunoblots of LAMP1, LC3 (LC3-I and LC3-II), and tau in human iPSC-derived neurons expressing EGFP control, UBA5, or UFM1 after 21 days in culture. (B, C) Quantification of LC3-I and LC3-II (B) and LAMP1 (∼100 kDa) (C) protein levels in neurons described in (A), demonstrating reduced lysosomal and autophagic markers upon UBA5 or UFM1 overexpression. (D) Quantification of high-molecular-weight (HMW) tau species (>70 kDa) in neurons described in (A), showing increased tau accumulation with UBA5 or UFM1 overexpression. n = 8 biological replicates from two independent experiments. Statistical significance was assessed by one-way ANOVA with Tukey’s multiple comparisons test (**p < 0.01, ***p < 0.001, ****p < 0.0001). (E, F) Confocal fluorescence microscopy (E) and quantitative analysis (F) of lysosomal parameters in iPSC-derived neurons expressing EGFP, UBA5-EGFP, or UFM1-EGFP. Scale bars, 20 µm. (G, H) Confocal fluorescence microscopy (G) and quantification (H) of MC1-positive tau inclusions in iPSC-derived neurons expressing EGFP, UBA5-EGFP, or UFM1-EGFP, with or without K18 tau seeding. Quantification represents the percentage of MC1-positive neurons per group. n = 12 biological replicates from two independent experiments. Statistical significance was determined by one-way ANOVA with Tukey’s multiple comparisons test (***p < 0.001, ****p < 0.0001). (I) Representative immunoblots of TRAPPC8, LAMP1, and LC3 in human iPSC-derived neurons expressing shRNA control or shRNA targeting TRAPPC8 after 21 days in culture. (J–L) Quantification of TRAPPC8 (J), LC3-I and LC3-II (K), and LAMP1 (L) in neurons described in (I), demonstrating lysosomal and autophagic impairment upon TRAPPC8 depletion. n = 8 biological replicates from two independent experiments. Statistical significance was assessed by unpaired two-tailed Student’s t test (****p < 0.0001). (M) Confocal fluorescence microscopy of MC1-positive tau aggregates in iPSC-derived neurons expressing shRNA control or shRNA targeting TRAPPC8. Scale bars, 20 µm. (N) Quantification of the percentage of MC1-positive neurons in (M). Statistical significance was determined by unpaired two-tailed Student’s t test (****p < 0.0001).

### The TRAPPC8 ASH domain binds UBA5 to suppress UFMylation

Our data so far showed TRAPPC8’s association with both UBA5 and UFM1 and its functions as a brake on UFMylation. To define the structural basis of TRAPPC8’s interaction with UBA5, we generated a series of truncation constructs spanning the full-length protein (**Fig. 5A**). Co-immunoprecipitation assays in HEK cells co-expressing UBA5-EGFP and either full-length or truncated TRAPPC8-V5 constructs demonstrated robust interaction between UBA5 and full-length TRAPPC8 (**Fig. 5B**). Domain-mapping analysis revealed that the 701–1100 fragment exhibited the strongest association with UBA5 (**Fig. 5C**), identifying the ASH domain–containing region as the principal interface mediating TRAPPC8–UBA5 interaction.

We next performed AlphaFold3 modeling of a complex comprising UBA5, mature UFM1, and the full length of TRAPPC8 (high light 701–1100 fragment) (**Fig. 5D–E**). The model predicts that a C-terminal α-helix of UBA5 engages the surface of the TRAPPC8 U5BR fragment. Notably, this α-helix overlaps with the previously defined UFC1-binding sequence (UBS) of UBA5 (*37*), suggesting that TRAPPC8 binding may sterically hinder UFC1 recruitment and thereby limit UFM1 transfer.

We directly test this hypothesis using a cell-free NanoBiT-based UFMylation assay that monitors UFM1 transfer from UBA5 to UFC1 via luminescence reconstitution (**Fig. 5F**). Recombinant UBA5, UFC1, and UFM1 generated robust luminescence, reflecting efficient UFM1 transfer (**Fig. 5G**). Addition of the TRAPPC8 U5BR fragment suppressed this signal in a dose-dependent manner. ATP omission or use of a catalytically inactive UBA5 C250A mutant abolished luminescence, confirming assay specificity (**Fig. 5G**). Thus, TRAPPC8 can directly inhibit UBA5-mediated UFM1 transfer under cell-free condition.

We then expressed V5-tagged TRAPPC8 (701–1100) in HEK cells and stimulated UFMylation with anisomycin (**Fig. 5H**). Expression of the U5BR fragment markedly reduced UFM1-conjugated species across multiple molecular weight ranges compared to controls (**Fig. 5H**). Quantification confirmed a significant suppression of UFM1 conjugation (**Fig. 5I–K**), indicating that this region is sufficient to inhibit UFMylation in a cellular context. Our results identify the ASH domain–containing 701–1100 region of TRAPPC8 as the principal UBA5-binding interface that restrains UFMylation, likely by limiting UBA5–UFC1 engagement.

**Figure 5.**
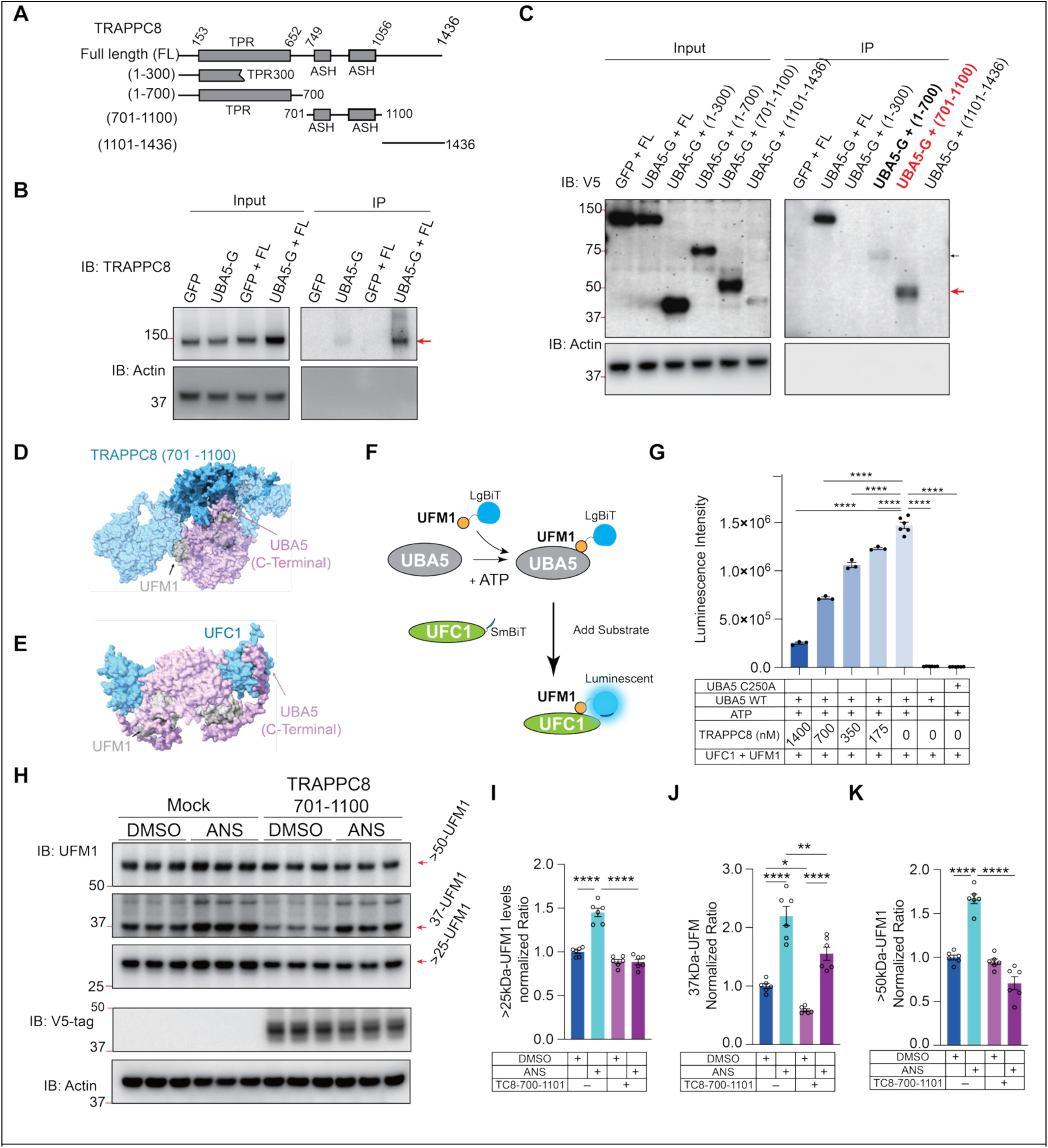
The TRAPPC8 ASH domain binds UBA5 to suppress UFMylation. (A) Schematic representation of full-length and truncated TRAPPC8 constructs used for domain-mapping analyses. The 701–1100 amino acid region contains duplicated ASH (ASPM-SPD-2-Hydin) domains and is designated the UBA5-binding region (U5BR). (B, C) Co-immunoprecipitation assays in HEK cells co-transfected with UBA5-EGFP and either full-length TRAPPC8-V5 (B) or truncated TRAPPC8-V5 constructs (C). Cell lysates were immunoprecipitated with anti-EGFP antibody and immunoblotted with anti-TRAPPC8 (B) or anti-V5 (C) antibodies to assess interaction with UBA5. (D, E) AlphaFold3-predicted structural models of the UBA5–UFM1–TRAPPC8 (Full length, high light aa 701–1100) complex (D) and the UBA5–UFM1–UFC1 complex (E), highlighting the overlapping interaction interface on UBA5-C-terminal. (F) Schematic of the cell-free NanoBiT-based UFMylation assay used to monitor UFM1 transfer from UBA5 to UFC1 through luminescence reconstitution. (G) Quantification of luminescence intensity in the NanoBiT assay. UFC1-SmBiT + LgBiT-UFM1 + UBA5 (WT) with ATP served as the positive control; UBA5 C250A mutant or ATP-free conditions served as negative controls. Data represent n = 3 biological replicates from two independent experiments. Statistical significance was determined by one-way ANOVA with Tukey’s multiple comparisons test (****p < 0.0001). (H) Representative immunoblots of UFM1, V5-tagged TRAPPC8 (701–1100), and β-actin in HEK cells transfected with control (mock) or TRAPPC8 (701–1100) expression plasmids. Cells were treated with 200 nM anisomycin for 10 min prior to harvest. (I–K) Quantification of UFM1-conjugated species (>25 kDa, ∼37 kDa, and >50 kDa) in HEK cells described in (H). n = 6 biological replicates from two independent experiments. Statistical significance was assessed by one-way ANOVA with Tukey’s multiple comparisons test (****p < 0.0001).

### TRAPPC8-ASH rescues UFMylation-induced lysosomal deficiency and tau propagation via restoring CLN8

We next determined if the ASH domain of TRAPPC8 is sufficient to restore lysosomal defects induced by hyper-UFMylation. In HEK cells expressing UFMylation constructs, CLN8 and LAMP1 levels were reduced (**Fig. 2** and **Fig. 4**), consistent with impaired lysosomal homeostasis. Co-expression of TRAPPC8 (701–1100) restored CLN8 and LAMP1 protein levels (**Fig. 6A–C**), demonstrating the functional competence of ASH domain of TRAPPC8 in restraining hyper-UFMylation-induced lysosomal dysfunction.

To determine whether CLN8 itself is sufficient to counteract UFMylation-induced tau aggregation, we overexpressed CLN8 in human iPSC-derived neurons expressing UFM1. Immunoblotting confirmed restoration of CLN8 levels (**Supplementary Fig. 5A** and **B**). Importantly, CLN8 overexpression significantly reduced MC1-positive tau inclusions and aggregate burden, indicating that CLN8 acts downstream of hyper-UFMylation to suppress Tau pathology (**Fig. 6D** and **6E**).

We next asked whether restoring the ASH domain of TRAPPC8 is sufficient to mitigate hyperufmylation-induced tau propagation. Human iPSC-derived neurons were transduced with EGFP control, UFM1, or UFM1 together with TRAPPC8 (701–1100) and maintained for 21 days. Immunoblot analysis confirmed robust expression of the V5-tagged TRAPPC8 fragment without altering endogenous full-length TRAPPC8 levels (**Supplementary Fig. 5C–E**). As expected, UFM1 overexpression markedly increased MC1-positive inclusions compared to control neurons. Co-expression of TRAPPC8 (701–1100) significantly reduced both the percentage of MC1-positive neurons and the somatic burden of aggregated tau (**Fig. 6F** and **6G**), indicating that ASH domain of TRAPPC8 is sufficient to mitigate tau pathology.

### The UFMylation-TRAPPC8 axis regulates Tau propagation in vivo and is disrupted in Alzheimer’s disease

To determine whether the UFMylation–TRAPPC8 axis regulates tau propagation in vivo, we used a unilateral fibril-seeding model in PS19 mice (**Fig. 6H**). K18 tau fibrils were injected into one hippocampus to initiate tau seeding, while the contralateral hippocampus received lentivirus expressing EGFP, UBA5, or the TRAPPC8 ASH domain (701–1100). Tau propagation was quantified by measuring MC1 immunoreactivity in the contralateral hippocampus normalized to the ipsilateral injection site. Compared with EGFP controls, UBA5 overexpression significantly enhanced contralateral tau propagation, indicating that hyper-UFMylation promotes tau spreading in vivo. In contrast, expression of the TRAPPC8 ASH domain markedly reduced tau propagation, demonstrating that restoration of TRAPPC8 activity is sufficient to suppress pathological tau spread in vivo (**Fig. 6I and 6J**).

To determine the relevance of this regulatory axis in human, we next examined TRAPPC8 levels in an independent cohort of postmortem AD brains obtained from the University of Pennsylvania Brain Bank. Neurons with pathological tau aggregates were identified by MC1 co-immunostaining (**Fig. 6K**). Among MC1-negative neurons, TRAPPC8 immunoreactivity was significantly reduced in AD brains compared to non-demented controls. Within AD brains, MC1-positive neurons exhibited a further decrease in TRAPPC8 levels relative to MC1-negative neurons (**Fig. 6L**). Together, these findings indicate that reduced TRAPPC8 levels are associated with increased tau aggregation and support a model in which TRAPPC8 functions as a negative regulator of tau pathology in AD.

**Figure 6.**
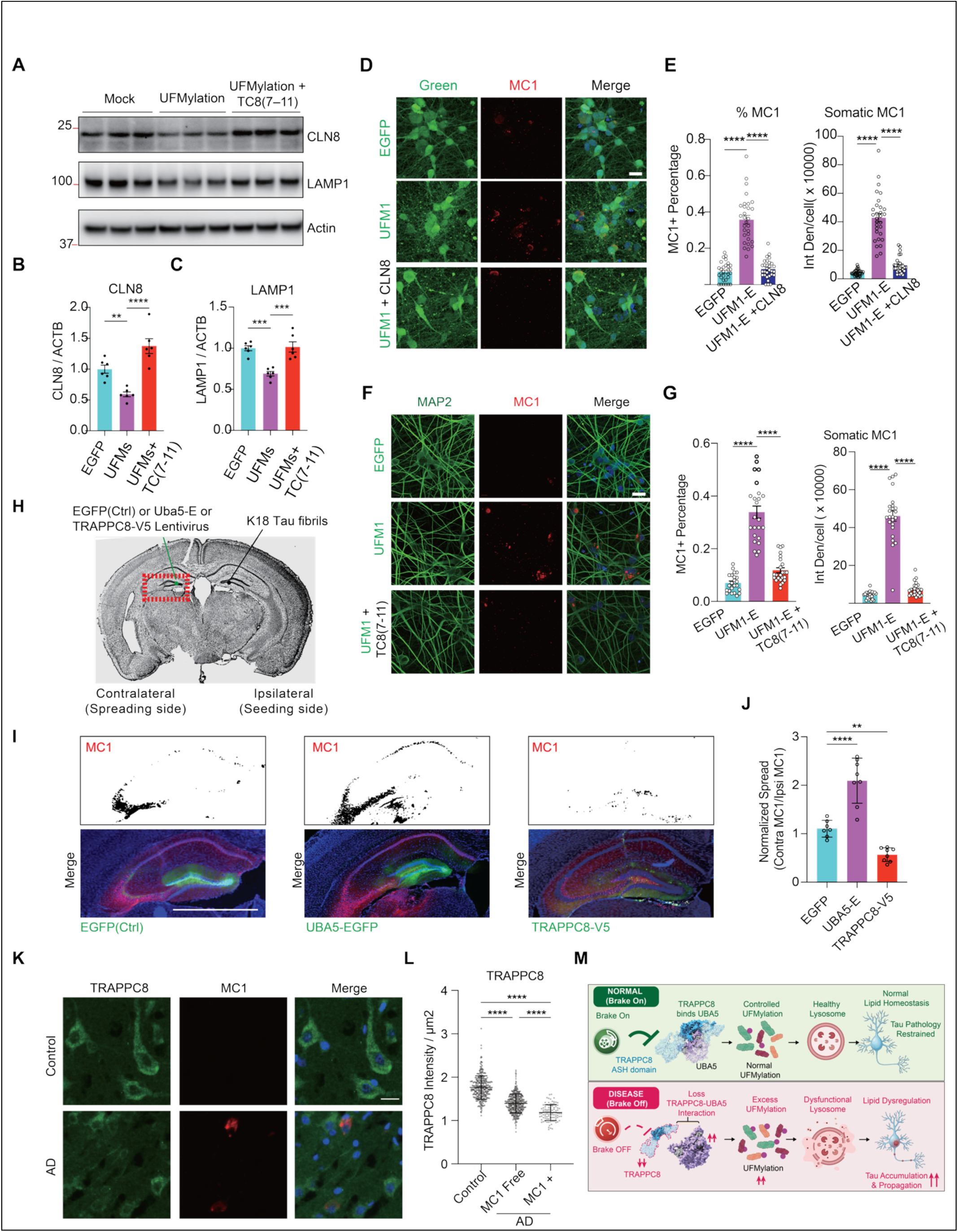
Restoration of UBA5-binding TRAPPC8 domain or CLN8 Rescues UFMylation-Induced Tau Pathology. (A) Representative immunoblots of CLN8, LAMP1, and β-actin in HEK cells transfected with control (mock), UFMylation constructs, or UFMylation constructs plus TRAPPC8 (701–1100). (B, C) Quantification of CLN8 (<25 kDa) (B) and LAMP1 (∼100 kDa) (C) protein levels in cells described in (F). n = 3 biological replicates. Statistical significance was determined by one-way ANOVA with Tukey’s multiple comparisons test (****p < 0.0001). (D) Confocal fluorescence microscopy of MC1-positive tau aggregates in iPSC-derived neurons expressing EGFP, UFM1, or UFM1 plus CLN8. Scale bars, 20 µm. (E) Quantification of the percentage of MC1-positive neurons and somatic integrated MC1 fluorescence intensity per cell in (D). n = 30 biological replicates from three independent experiments. Statistical significance was assessed by one-way ANOVA with Tukey’s multiple comparisons test (****p < 0.0001). (F) Confocal fluorescence microscopy of MC1-positive tau aggregates in iPSC-derived neurons expressing EGFP, UFM1, or UFM1 plus TRAPPC8 (701–1100). Scale bars, 20 µm. (G) Quantification of the percentage of MC1-positive neurons and somatic integrated MC1 fluorescence intensity per cell in (F). n = 24 biological replicates from three independent experiments. Statistical significance was assessed by one-way ANOVA with Tukey’s multiple comparisons test (****p < 0.0001). (H) Diagram of PS19 mouse brain indicating lentivirus (spreading) and K18-Tau fibril (seeding) injection sites. (I) Representative images of MC1 immunostaining in the mouse brain after control(EGFP) or UBA5, or TRAPPC8 overexpression lentivirus injections. Scale bar, 2000 μm. (J) Quantification of relative MC1+ signal from the hippocampus of individual groups from (I), normalizing the spreading side to the seeding side. n = 23 EGFP, 28 UBA5-EGFP and 27 TRAPPC8-V5 replicates in 7 EGFP, 8 UBA5-EGFP and 8 TRAPPC8-V5 mice. ∗∗p < 0.01, ****p<0.0001, mixed model analysis. (K) Representative immunofluorescence images of human AD postmortem tissue stained with TRAPPC8, MC1 antibodies and DAPI. Samples are from the University of Pennsylvania (UPenn) Brain Bank. Scale bar, 20 μm. (L) Quantification of TRAPPC8 immunofluorescence normalized individual cell area µm^2^ in human AD postmortem tissue. n = 299 from control, n = 381 from MC1free cells and 98 MC1+ cells from four controls, and six pathological cases. ∗∗∗∗p < 0.0001, mixed model analysis. (M) Schematic model illustrating UFMylation-TRAPPC8-CLN8 axis linking ubiquitin-like regulation, lysosomal homeostasis, and tau propagation in neurodegenerative disease

## Discussion

The strong correlation between pathological tau burden and cognitive decline in Alzheimer’s disease (AD) has motivated the search for upstream proteostatic mechanisms that govern tau aggregation and propagation (*37–39*). Our findings identify TRAPPC8 as the endogenous homeostatic brake that restrains neuronal UFMylation through direct regulation of the E1 enzyme UBA5, thereby defining a previously unrecognized mechanism that sets the activity of this ubiquitin-like modification pathway. Loss of this regulatory mechanism likely contributes to sustained UFMylation hyperactivation, lysosomal dysfunction, lipid dyshomeostasis, and tau pathology in AD. Importantly, restoration of the TRAPPC8 UBA5-binding region is sufficient to suppress tau pathology in vivo, demonstrating that re-establishing endogenous UFMylation homeostasis represents a potential therapeutic strategy for tauopathies (**Fig. 6M**).

Although the enzymatic cascade responsible for UFMylation has been well established, including the E1 enzyme UBA5, the E2 enzyme UFC1, the E3 ligase UFL1, and the de-UFMylating proteases UFSP1 and UFSP2, how neurons physiologically regulate overall UFMylation activity has remained unclear (*40–42*). Our study fills this conceptual gap by identifying TRAPPC8 as an endogenous regulator that limits UFMylation upstream of substrate modification. Unlike UFSP1/2, which reverse UFMylation after conjugation, TRAPPC8 restricts pathway activity before UFM1 is transferred to UFC1, representing a fundamentally distinct mode of regulation. This suggests that adaptor-mediated control of E1 activity may constitute a broader regulatory principle within ubiquitin-like modification systems.

Consistent with this model, AD cortex exhibits elevated high-molecular-weight UFM1 conjugates without a corresponding increase in free UFM1, whereas TRAPPC8 immunoreactivity is markedly reduced in MC1-positive tau-bearing neurons, supporting enhanced UFMylation activity rather than increased UFM1 expression. Together with previous reports describing reduced UFSP2 expression in AD (*13, 14*), these findings indicate that disruption of multiple regulatory mechanisms may converge to drive pathological UFMylation hyperactivation during disease progression. Importantly, the bidirectional regulation between TRAPPC8 depletion and UFMylation hyperactivation therefore creates a self-reinforcing circuit that progressively destabilizes neuronal proteostasis.

Furthermore, our findings identify CLN8 as a critical downstream effector of the TRAPPC8–UFMylation pathway. Suppression of CLN8 by hyper-UFMylation provides a mechanistic link between elevated UFMylation and lysosomal dysfunction, whereas restoration of CLN8 is sufficient to attenuate tau pathology. Consistent with the emerging role of CLN8 in lipid homeostasis, hyper-UFMylation also caused accumulation of neutral lipids and lipid droplets, suggesting that defective lysosomal trafficking and lipid homeostasis constitute key downstream mechanism linking UFMylation hyperactivation to tau pathology (*29, 43*).

From a translational perspective, expression of the UBA5-binding region of TRAPPC8 was sufficient to suppress tau pathology in vivo, demonstrating that enhancing endogenous restraint of UFMylation can ameliorate disease-associated pathology. These findings suggest that small molecules, stabilized peptides, or gene therapy approaches designed to enhance the TRAPPC8–UBA5 interaction may represent promising therapeutic strategies. Alternatively, strategies to augment CLN8 activity or lysosomal enzyme trafficking may prove beneficial in the context of UFMylation-driven tau pathology. These approaches are complementary to ongoing efforts to target the UFMylation cascade pharmacologically and to emerging interest in lysosome-directed therapies for tauopathies.

Our proteomic analyses further identify MYH9 as a direct neuronal substrate of UFMylation through modification of Lys1387. Given the central role of MYH9 in cytoskeletal organization and vesicular trafficking (*44*), its modification may represent an additional mechanism through which UFMylation influences neuronal proteostasis and tau propagation. Defining how UFMylation at Lys1387 regulates MYH9 function and neuronal homeostasis will be an important direction for future studies. Although important questions remain—including the precise neuronal UFMylome, the functional consequences of MYH9 UFMylation, and the role of aging and non-neuronal cells—our findings establish a conceptual framework for understanding how neuronal UFMylation is physiologically restrained and how disruption of this homeostatic mechanism contributes to tauopathy.

## Acknowledgments

We would like to thank Guillermo Coronas and Ana Carney for administrative support, Weill Cornell Medicine’s Proteomics & Metabolomics Core, and Microscopy and Imaging Analysis Core. We thank Dr. Pingwei Li at Texas ACM University for the gift of the Ulp1 SUMO protease plasmid. We also thank Xiaoman Wang for helpful dry lab technical assistance and data analysis.

## Author contributions

L.G., and Y.W., conceptualized the study. Y.W., L.G., H.L., and H.Y.Y., designed experiments. Y.W. performed majority of the experiments and data analysis in this paper. E.C.performed iPSC neuron culture, immunocytochemistry and WB under the supervision of Y.W.. S.L., W.F., J.Z., H.C., B.G., R.S., assistanted iPSC culture and WB. M.Y.W. assistanted human brain IF. H. C., and B. L. assistanted proteomic and lipidomic data analysis. Y.W., S.W., B.G., and S.G. designed and made plasmids. S.A. organized human brain postmortem tissues from the Mount Sinai brain bank. K.M. performed GFP-IP-MS and Y.S. conducted data analysis under the supervision of H.Y. with assistance from X.W.. Z.G. perfomed protein purification and inhibitory assay under the supervision of Z.L. and H.L.. Y.W. and L.G. wrote the manuscript. L.G. supervised the study.

## Supplementary Materials

Materials and Methods

Figs. S1 to S6

Tables S1 to S6

